# Exercise prevents cardiac electrical remodeling in doxorubicin-treated female mice but does not provide cardioprotection in males

**DOI:** 10.64898/2026.05.28.728478

**Authors:** Amber V Melcher, Lauren Hafflett, Lichao Tang, Katy Trampel, Manas Bodapotula, Sharon Ann George

## Abstract

**Background:** Doxorubicin (DOX) causes sex-specific cardiotoxicity. Metabolic impairment is a well-established cardiotoxic effect of DOX treatment that can contribute to other detrimental effects such as increased reactive oxygen species, reduced ATP, inflammation etc. We hypothesized that preserving cardiac metabolism by exercise can attenuate DOX cardiotoxicity.

**Methods:** Male and female C57BL/6J mice at 15 weeks of age were randomly assigned to one of four groups, 1) Control (sedentary), 2) EX (exercised, treadmill running), 3) DOX (doxorubicin at 5 mg/kg/week for 6 weeks), and 4) EXDOX (exercise + doxorubicin). Echocardiography was performed every other week during the 6-week protocol to measure cardiac mechanical function. At the end of the protocol, optical mapping and seahorse analysis were performed to measure electrophysiology and metabolism, respectively. RNA sequencing, cytokine array assay and transmission electron microscopy were also performed to determine sex-specific mechanisms of DOX cardiotoxicity.

**Results:** DOX reduced stroke volume and left ventricular diameter in males only and exercise did not prevent these effects of DOX. In female mice, DOX prolonged action potential duration (APD) and slowed conduction velocity (CV), and importantly, exercise prevented DOX-induced CV slowing. Exercise-induced cardioprotection against DOX in female mice was associated with preservation of aerobic metabolism and attenuation of inflammation which modulated ion channel gene expression. Specifically, Cacna1c was increased in both DOX and EXDOX females, but not in males and correlated with APD prolongation. Interestingly, despite CV slowing, Gja1 and Scn5a were increased. However, increased Kcnj8 along with metabolic impairment could cause membrane hyperpolarization and underlie CV slowing.

**Conclusions:** DOX cardiotoxicity is sex specific. Mechanical dysfunction is more prevalent in DOX-treated males while arrhythmogenic electrical remodeling is more prevalent in DOX-treated females. Exercise therapy during DOX did not prevent DOX induced mechanical dysfunction in male hearts but attenuated electrical remodeling in females by preserving metabolism and attenuating inflammation.

## INTRODUCTION

With advances in cancer therapies and improved survival among cancer patients, comes a new wave of healthcare problems, in the form of cancer therapy induced toxicity. Several clinically used chemotherapeutics have well-documented acute and chronic cardiotoxicity.^1^ Cardiotoxic effects are dose-dependent, cumulative and is also affected by the age and sex of the patient.^2–4^ Clinically, cardiotoxicity is primarily assessed by reduction in ejection fraction (EF). Reduction in EF to less than 50-55% or a greater than 10% decline since start of therapy is an indicator for re-evaluation of cancer therapy and treatment for cardiotoxicity.^5,6^ Current treatment of clinically diagnosed cardiotoxicity mostly follows heart failure guidelines with prescription of beta blockers, statins and ACE inhibitors.^7–9^

Doxorubicin (DOX, Adriamycin) is a widely used chemotherapeutic agent that induces double strand DNA breaks, activates topoisomerase 2B and disrupts cellular function by metabolic dysfunction, calcium mishandling, inflammation, stress signaling activation, autophagy dysregulation, and mitochondrial impairment, among other effects.^10,11^ It has been speculated that metabolic dysfunction is a primary cardiotoxic effect which can trigger or at least contribute to the activation of the other known downstream cardiotoxic effects of DOX.^12,13^ Metabolic dysfunction induced by DOX results in lower ATP production rate, substrate switching from fatty acids to glucose metabolism, lipid accumulation, and increased reactive oxygen species generation. In this study, we aimed to preserve metabolism during DOX therapy by aerobic exercise therapy.

Exercise therapy is now routinely prescribed for patients with cardiovascular diseases ranging from coronary heart disease to heart failure.^14,15^ Moderate endurance exercise improves cardiac metabolism, increases ATP production, mitigates oxidative stress and has anti-inflammatory effects.^16^ However, a large clinical study that aimed to assess the benefits of exercise therapy in heart failure patients have produced underwhelming results which were attributed to low adherence rate to prescribed exercise regimens among patients.^17^ We hypothesized that exercise prevents cardiac metabolic dysfunction during DOX therapy and can attenuate DOX induced cardiotoxicity.

In this study, we tested the benefits of exercise therapy in a chronic mouse model of DOX induced cardiotoxicity. We determined that not only is DOX cardiotoxicity sex dependent but cardioprotection with exercise therapy is also sex dependent. While exercise attenuated female DOX cardiotoxicity, it did not benefit males. These findings could have significant implications in clinical application of exercise therapy in cancer patients undergoing DOX chemotherapy and in cardiovascular diseases of similar etiologies.

## METHODS

All animal studies were approved by the Institutional Animal Care and Use Committee at Northwestern University and the University of Pittsburgh and were in accordance to the NIH’s Guide for Care and Use of Laboratory Animals.

*Mouse and Experimental Groups:* C57BL/6J mice (JAX, 000664) of both sexes at 15 weeks of age were used in this study. Mice were randomly divided into four groups (Figure 1A) 1) Control (sedentary), 2) EX (exercised), 3) DOX (administered DOX), and 4)

**Figure 1.**
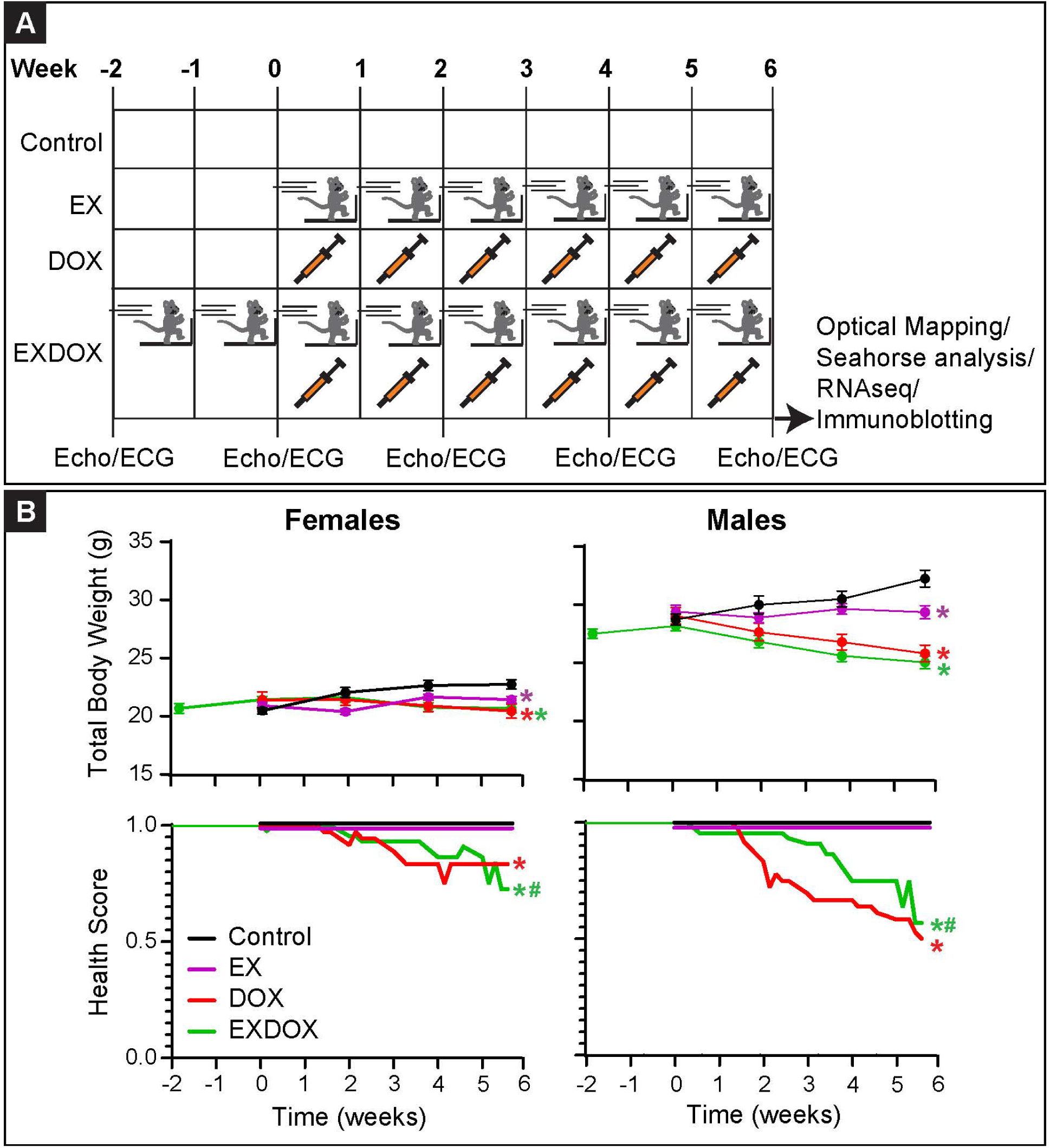
Experimental Design and DOX-induced morbidity. **A)** Mice were divided into four experimental groups 1) Control – sedentary mice, 2) EX – run on treadmill for 45 mins/day, 5 days/week for 6 weeks, 3) DOX – treated with 5 mg/kg/week intraperitoneal injections for 6 weeks, and 4) EXDOX – 2 weeks of EX alone followed by 6 weeks of EX + DOX. **B)** Change in total body weight over the study period in female (left) and male (right) mice in each of the four experimental groups is illustrated in the panels above and change in health score (decrease indicates morbidity) in the above groups is illustrated in the panels below. Sample sizes for both sexes in each experimental group are as follows: Control: 5/6 (males/females) mice, EX: 9 mice, DOX: 9 mice, EXDOX: 11 mice. Two-way ANOVA tests with time and experimental groups as the two variables were performed to determine significant differences between experimental groups. * indicates p<0.05 versus Control and # indicates p<0.05 versus DOX.

EXDOX (exercised and administered DOX). Mice were exercised by treadmill running at ∼15 m/min with no incline for 5 days/week. Mice in the EX group underwent exercise for 6 weeks. DOX was administered by weekly intraperitoneal injections at 5 mg/kg for 6 weeks. Mice in the EXDOX group were subjected to 2 weeks of exercise alone, followed by 6 weeks of DOX and exercise.

During the study period, mice were assessed 5 times a week and assigned a health score based on a previously published mouse health score rubric.^18,19^ Briefly, signs of distress such as hunched posture, ruffled fur, porphyrin accumulation around the eyes, slow or no movement in response to interaction, were assessed and a health score between 1 (healthy, no signs of distress) or 0 (dead) was assigned. Body weight was also determined every 2 weeks.

*Echocardiography:* At the start of the study period (Week 0) and every 2 weeks thereafter, left ventricular (LV) structure and mechanical function were assessed by echocardiography. Mice were anesthetized using an isoflurane vaporizer (1-2.5% isoflurane in 1 ml/min oxygen) and transferred to the heated platform of a Vevo 3100 Ultrasound Machine (Visualsonics). Chest hair was removed from the sternum to the left side of the mouse and ultrasound gel (Aquasonics, Clear) was applied to this region. Transthoracic M-mode echocardiography was performed, and data were analyzed using the Vevo LAB 5.8.2 software. Ejection Fraction (EF), Fractional Shortening (FS), Stroke Volume (SV), Cardiac Output (CO), and LV End Diastolic/Systolic Volumes (LVEDD/LVESD) were measured from the echocardiograms.

*Triple Parametric Optical Mapping:* At the end of the 6-week study period, triple parametric optical mapping to measure cardiac metabolism-excitation-contraction coupling was performed.^20^ Mice were euthanized by isoflurane inhalation followed by cervical dislocation and thoracotomy for heart excision. The aorta was cannulated and the heart was retrogradely perfused with a modified Tyrode’s solution (in mM, 130 sodium chloride, 24 sodium bicarbonate, 1.2 monosodium phosphate, 1 magnesium chloride, 5.6 glucose, 4 potassium chloride and 1.8 calcium chloride, pH at 7.4 and temperature maintained at 37°C) using a Langendorff perfusion system and immersed in a bath containing the same perfusate. Perfusion pressure was maintained at ∼80 mmHg by adjusting the perfusate flow rate.

After 10 mins of acclimatization, RH237 (30 μl of 1.25 mg/ml stock dye solution diluted in perfusate to 1 ml) and then Rhod-2 AM (30 μl of 1 mg/ml stock dye solution diluted in 30 μl Pluronic F-127 and perfusate to 1 ml) were slowly injected into the system through a dye injection port immediately above the cannula, with a 5 min washout period after each dye injection. The heart was paced by a platinum bipolar electrode placed on the anterior surface of the heart using stimuli at 2 ms duration, amplitude at 1.5x the threshold of stimulation and at basic cycle lengths (BCLs) ranging from 200 ms up to loss of 1:1 capture or arrhythmia. Hearts were illuminated with blue (365 nm) and green (520 nm) high powered LEDs light sources (Prizmatix) to induce NADH autofluorescence and to excite RH237 and Rhod-2 AM, respectively. Emitted light was recorded by a triple parametric optical mapping system in tandem lens configuration where the three cameras recorded NADH, RH237 and Rhod-2 AM signals simultaneously, from the same field of view.

From the recorded optical signals, ten parameters of cardiac function were measured using a Matlab based data analysis software, Rhythm 3.0.^20^ From transmembrane potential optical recordings, voltage rise time (V_m_ RT): time interval of 10 to 90% of the upstroke of the action potential, action potential duration (APD_80_): interval between activation time defined as dF/dt_max_ and 80% repolarization, transverse and longitudinal conduction velocities (CV_T_ and CV_L_, respectively): speed of electrical wavefront propagation in the defined directions, and anisotropic ratio (AR): ratio of CV_L_/CV_T_ were calculated. From the intracellular calcium optical recordings, calcium rise time (CaRT): interval from 10 to 90% of calcium transient upstroke, calcium transient duration (CaTD_80_): interval from dF/dt_max_ in the calcium upstroke/release phase to 80% calcium reuptake, and calcium decay constant (Ca τ): time constant of exponential fit to the calcium reuptake phase of the transient were calculated. Additionally, voltage calcium delay (V_m_-Ca delay) was calculated as the interval between activation defined as dF/dt_max_ in transmembrane potential trace to it corresponding calcium transient. Lastly, absolute NADH intensity was determined for each condition and normalized to the first recording for each given heart to account for interexperimental variability.

*Seahorse Metabolic Analysis:* At the end of the six-week study period, hearts from one set of mice were excised and immediately arrested by perfusion and submersion in ice-cold cardioplegic solution (in mM: 110 NaCl, 16 KCl, 16 MgCl2, 1.5 CaCl2, and 10 NaHCO3 at pH 7.4). The left ventricle was then isolated and attached to the tissue holder of a precision vibrating microtome and continued to be submerged in cardioplegia. Slices were generated using the following setting: thickness of 100 μm, amplitude of 1 mm, frequency of 80 Hz and speed of 0.04 mm/s. Slices were transferred to a separate dish which contained the same cardioplegic solution and kept flat by placing a plastic mesh over it. Once all the slices were collected, slices were attached to the bottom of a Seahorse microculture plate (Agilent) using CellTak (Corning) as per manufacturer’s instructions. Seahorse XF DMEM media supplemented with 0.61 mM BSA-palmitate, 10 mM glucose, 1 mM pyruvate and 2 mM glutamine was prepared and added to each well in the microculture plate, careful not to agitate the slices and cause then to detach from the plate. Oxygen consumption rate (OCR) and extracellular acidification rate (ECAR) were measured using a Seahorse XFe96 analyzer for each experimental group, as previously described.^21^

*RNA sequencing:* At the end of the optical mapping studies, hearts were fixed in RNAlater and RNA sequencing was performed at Azenta Life Sciences as previously described.^22^ Differentially expressed genes (DEGs) were determined in R software using the DESeq2 package, with significance level set to <0.05. PCA plots, volcano plots and heat maps were generated from DEGs. Gene set enrichment analysis was performed using clusterProfiler package and gene sets enriched in Control versus experimental group is indicated in black while gene sets enriched in an experimental group versus control is indicated by corresponding group’s color.

*Cytokine Array:* At the end of the 6-week study protocol, one set of hearts were frozen and tissue lysates were generated in RIPA lysis buffer supplemented with protease and phosphatase inhibitors. Pooled samples from female hearts were used to determine the protein expression of 40 pro- and anti-inflammatory cytokines/chemokines using the Mouse Inflammation Array (RayBiotech, Cat # AAM-INF-1) by following the manufacturer’s instructions. Expression of each cytokine/chemokine was determined by measuring the spot density associated with each target using ImageJ and normalized to the controls on the array as indicated by manufacturer.

*Statistics:* Data analysis was performed using GraphPad Prism or R software (for RNA sequencing data analysis). All bars in the summary data represent group means and are superimposed with the individual data points. When present, error bars represent standard deviation unless noted otherwise in the figure legend. For data presented in Figure 1, two-way ANOVA tests were performed with experimental group as one variable and study time point as the second variable. For data presented in Figure 2, one-way ANOVA tests were first performed with experimental groups as the variable and unpaired, two-tailed Student’s t-tests were performed for post-hoc comparisons. For data presented in Figure 3, mixed effects model analysis with Geisser-Greenhouse correction was performed to determine significant differences in restitution curves between experimental groups. For data presented in Figure 4 and 6, unpaired, two-tailed Student’s t-tests were performed to determine significant differences between experimental groups. For data presented in Figure 5 and 7, Wald test was performed using DESeq2 package in R to determine statistically significant DEGs. Benjamini-Hochberg correction was applied for all multiple comparison corrections. p<0.05 is reported as statistically significant. * indicates p<0.05 vs Control and # indicates p<0.05 vs DOX.

**Figure 2.**
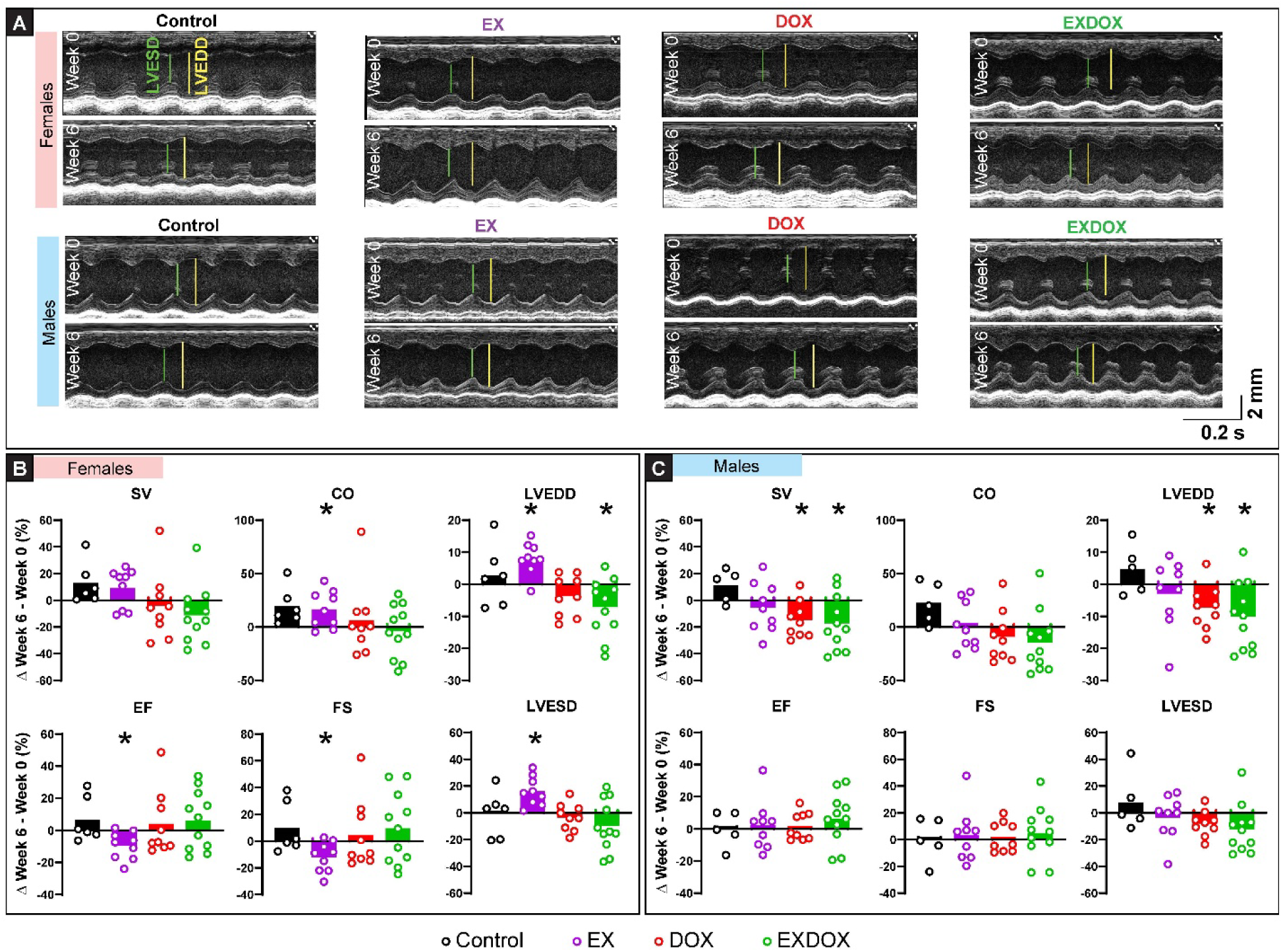
Exercise does not prevent DOX-induced mechanical dysfunction in male mice. **A)** Representative echocardiograms from male and female mice in the Control, EX, DOX and EXDOX groups. Summary data of echocardiographic analysis from female **(B)** and male **(C)** mice from the above four experimental groups. SV: stroke volume, EF: ejection fraction, CO: cardiac output, FS: fractional shortening, LVEDD: LV end diastolic diameter, LVESD: LV end systolic diameter. Sample sizes for both sexes in each of the experimental groups are as follows: Control: 5/6 (males/females), EX: 9, DOX: 9, EXDOX: 11. One-way ANOVA tests were performed to determine statistical differences between experimental groups followed by unpaired, two-tailed Student’s t-tests for post hoc analysis. Benjamini-Hochberg correction was applied for multiple comparisons. * indicates p< 0.05.

**Figure 3.**
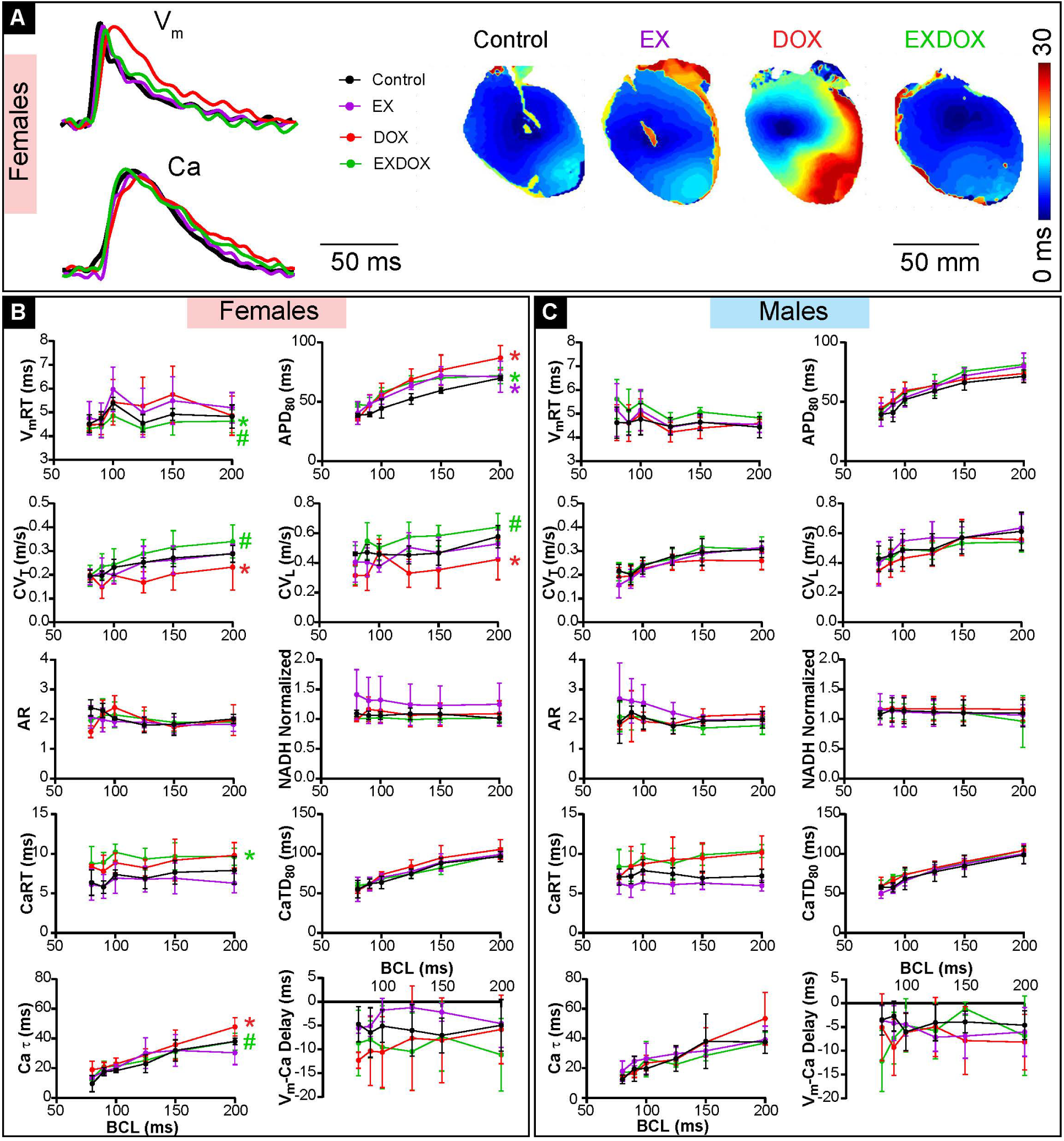
Exercise attenuates DOX-induced arrhythmogenic electrical remodeling in female mice. **A)** Representative action potential (top, left) and calcium transients (bottom, left) traces and activation maps illustration spread of electrical excitation over the mouse hearts surface in the four experimental groups, Control, EX, DOX and EXDOX. Summary data of ten parameters of cardiac function from female **(B)** and male **(C)** hearts recorded at BCLs ranging from 80 to 200 ms. VmRT: voltage rise time, APD80: action potential duration at 80% repolarization, CVT: transverse conduction velocity, CVL: longitudinal conduction velocity, AR: anisotropic ratio, CaRT: calcium rise time, CaTD80: calcium transient duration, Ca τ: calcium decay constant, Vm-Ca delay: voltage to calcium activation delay, NADH: normalized intensity of NADH autofluorescence. Sample sizes for both sexes in each experimental group are as follows: Control: 5, EX: 5, DOX: 6 and EXDOX: 6. Mixed-effects model analysis was performed with Geisser-Greenhouse correction to determine significant differences in restitution response in each of the parameters versus Control or DOX. * indicates p<0.05 versus Control and # indicates p<0.05 versus DOX.

**Figure 4.**
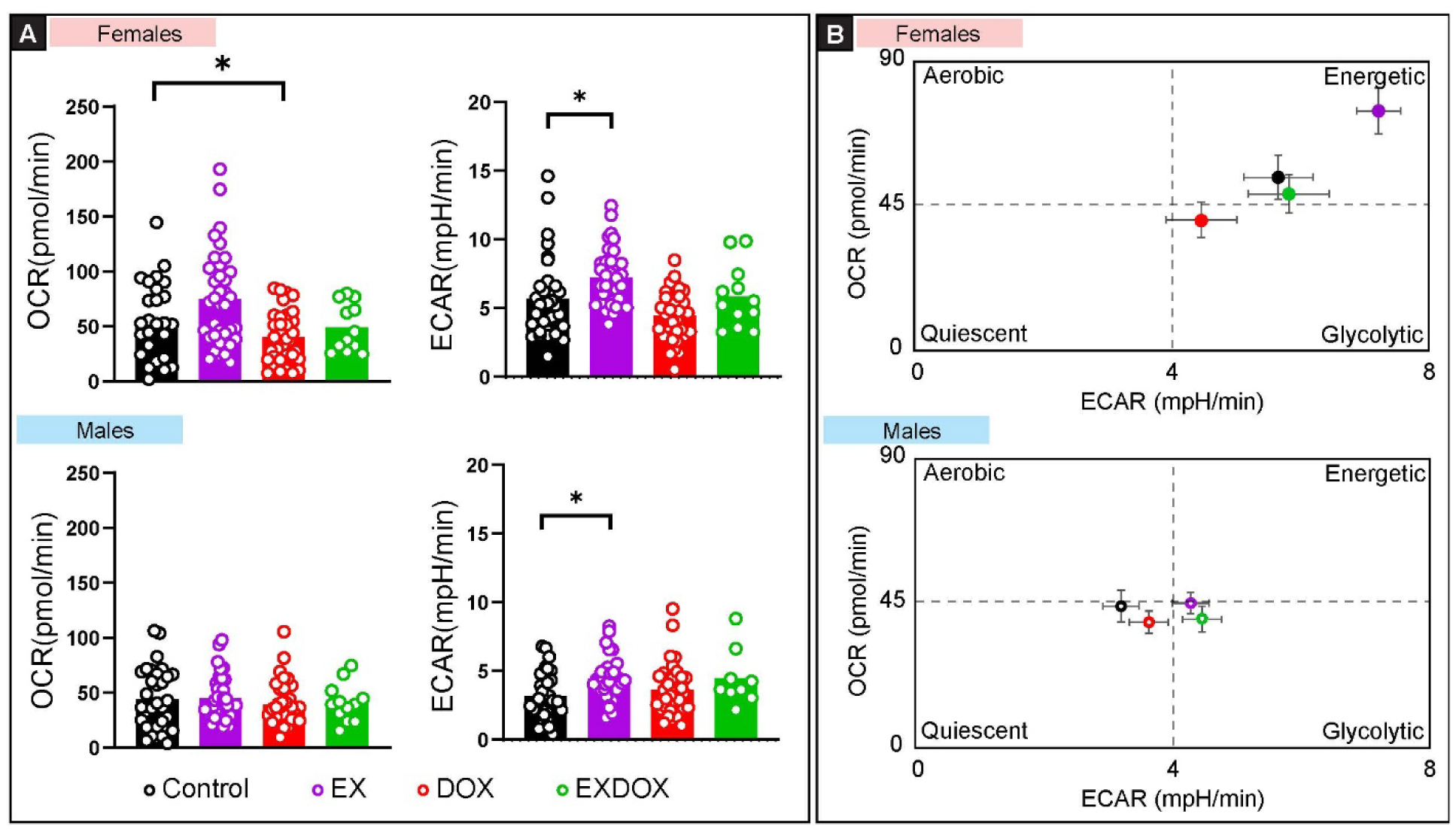
Exercise prevents DOX-induced metabolic impairment in female hearts. **A)** OCR (left) a measure of aerobic metabolism and ECAR (right) a measure of anaerobic metabolism was measured from male and female hearts I the four experimental groups, Control, EX, DOX and EXDOX. **B)** OCR vs ECAR values were plotted to generate the energy profile plots illustrating change in metabolic state under each condition. Error bars represent standard error. OCR: oxygen consumption rate, ECAR: extracellular acidification rate. Sample sizes: In each experimental group, 3 biological replicates were used, with 12 technical replicates from each heart. Unpaired, two-tailed Student’s t-tests were performed to determine statistical significance between experimental groups. Benjamini-Hochberg correction was applied for multiple comparisons. * indicates p<0.05 versus Control.

**Figure 5.**
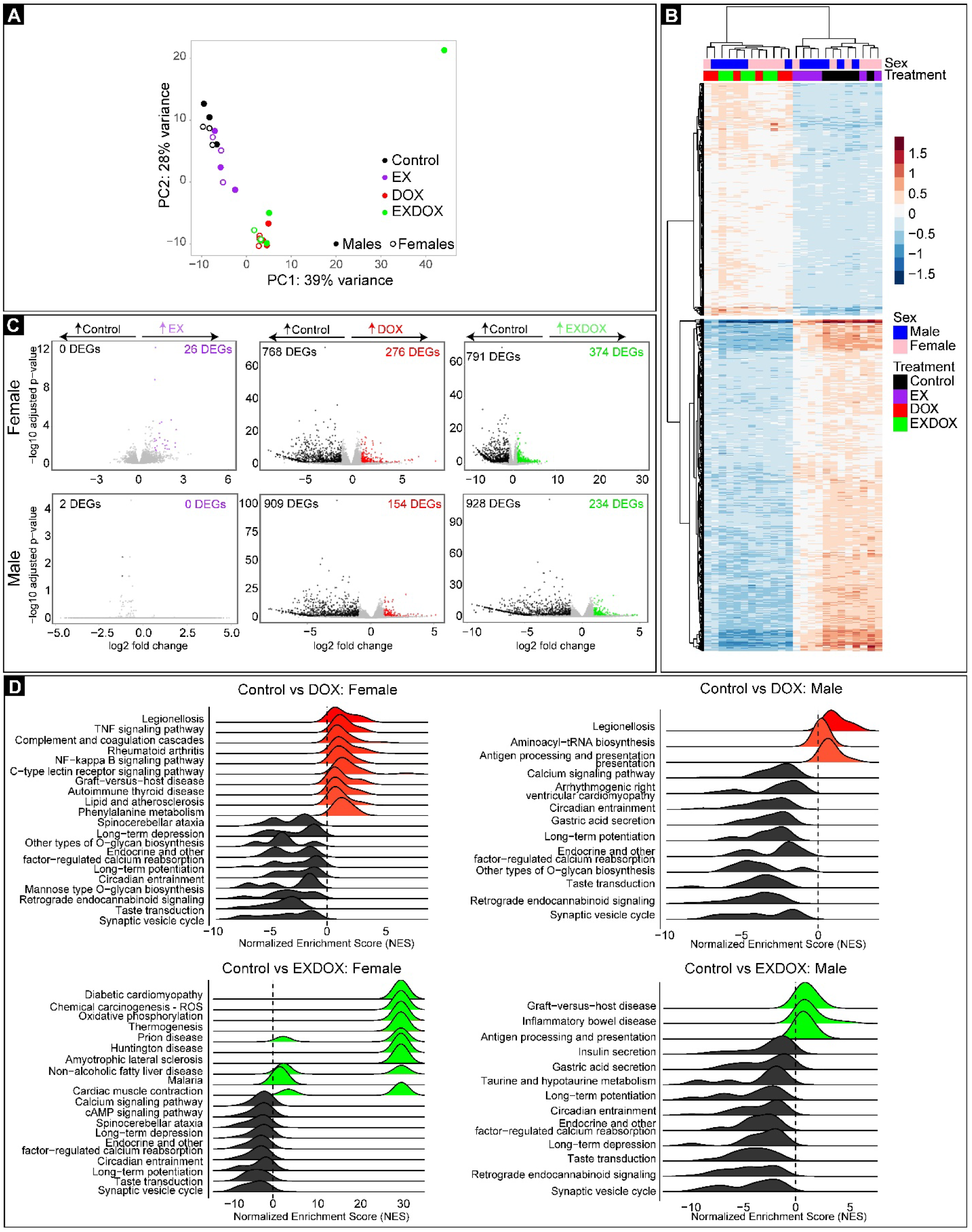
Sex-specific modulation of cardiac transcriptome by DOX. **A)** PCA plot illustrating clustering of experimental groups based on DOX treatment. **B)** Heat maps generated by uncharacterized hierarchical clustering analysis illustrating similarity in DOX treated groups of both sexes regardless of exercise. **C)** Volcano plots illustrating that while the cardiac transcriptome is not significantly altered by EX alone, DOX treatment with or without exercise generate a large number of DEGs. **D)** Top ten upregulated and downregulated gene sets in the male and female Control versus DOX and Control vs EXDOX groups. DEGs: differentially expressed genes. Sample sizes: In each experimental group, 3 biological replicates were used. DEGs were determined using the DESeq2 package in R which uses the Wald test to determine statistical significance and Benjamini-Hochberg method for multiple comparison correction.

**Figure 6.**
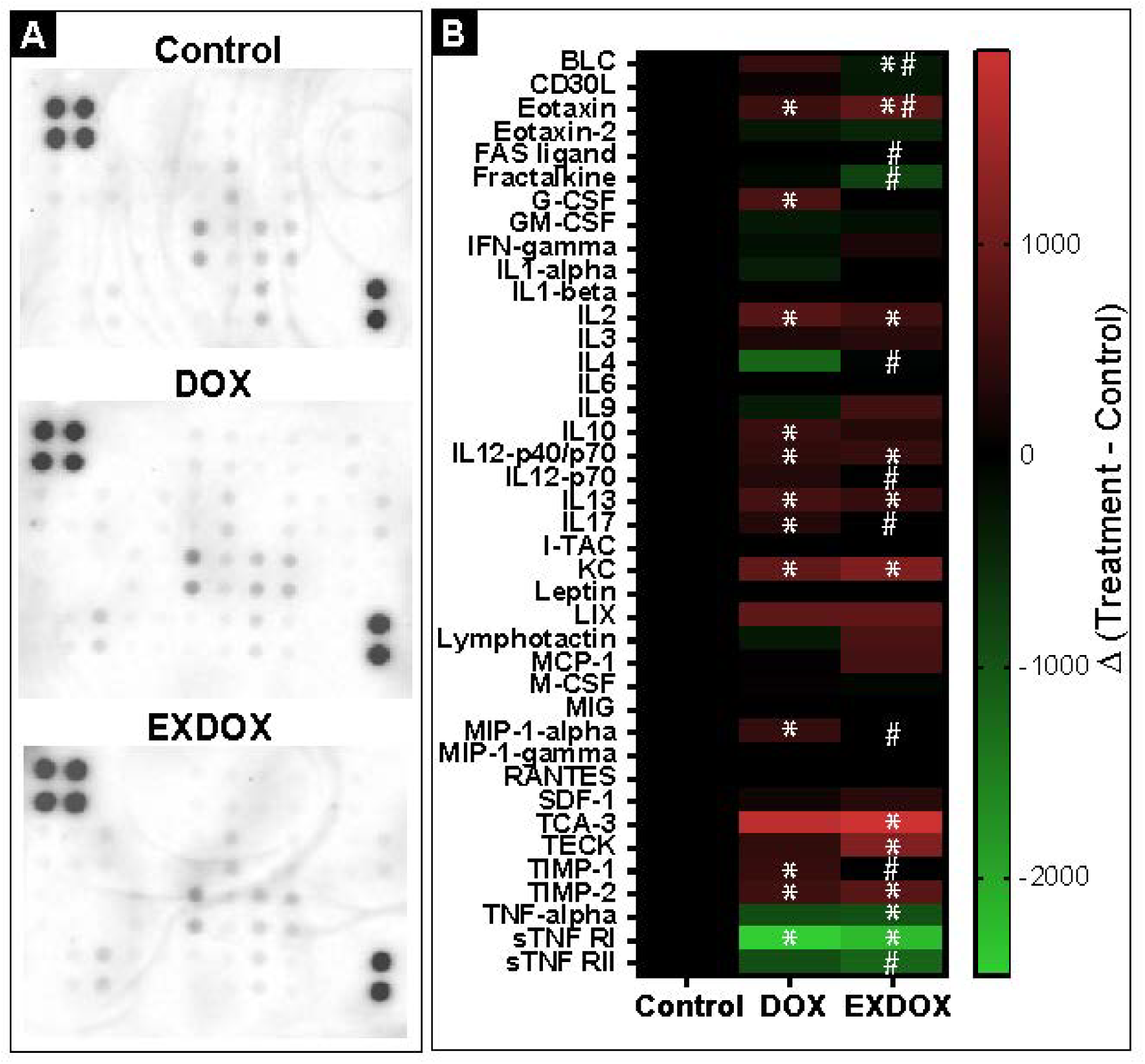
Exercise attenuates DOX-induced inflammatory condition in female hearts. **A)** Cytokine array blots illustrating modulation of 40 cytokines/chemokines in female Control, DOX and EXDOX groups. **B)** Heat map summarizing modulation of cytokine/chemokine protein expression in the above experimental groups. Sample sizes: Pooled samples of 3 biological replicates were used in each experimental group. Unpaired, two-tailed Student’s t-tests were performed to determine statistical differences between groups and Benjamini-Hochberg test was performed for multiple comparison correction. * indicates p<0.05 versus Control and # indicates p<0.05 versus DOX.

**Figure 7.**
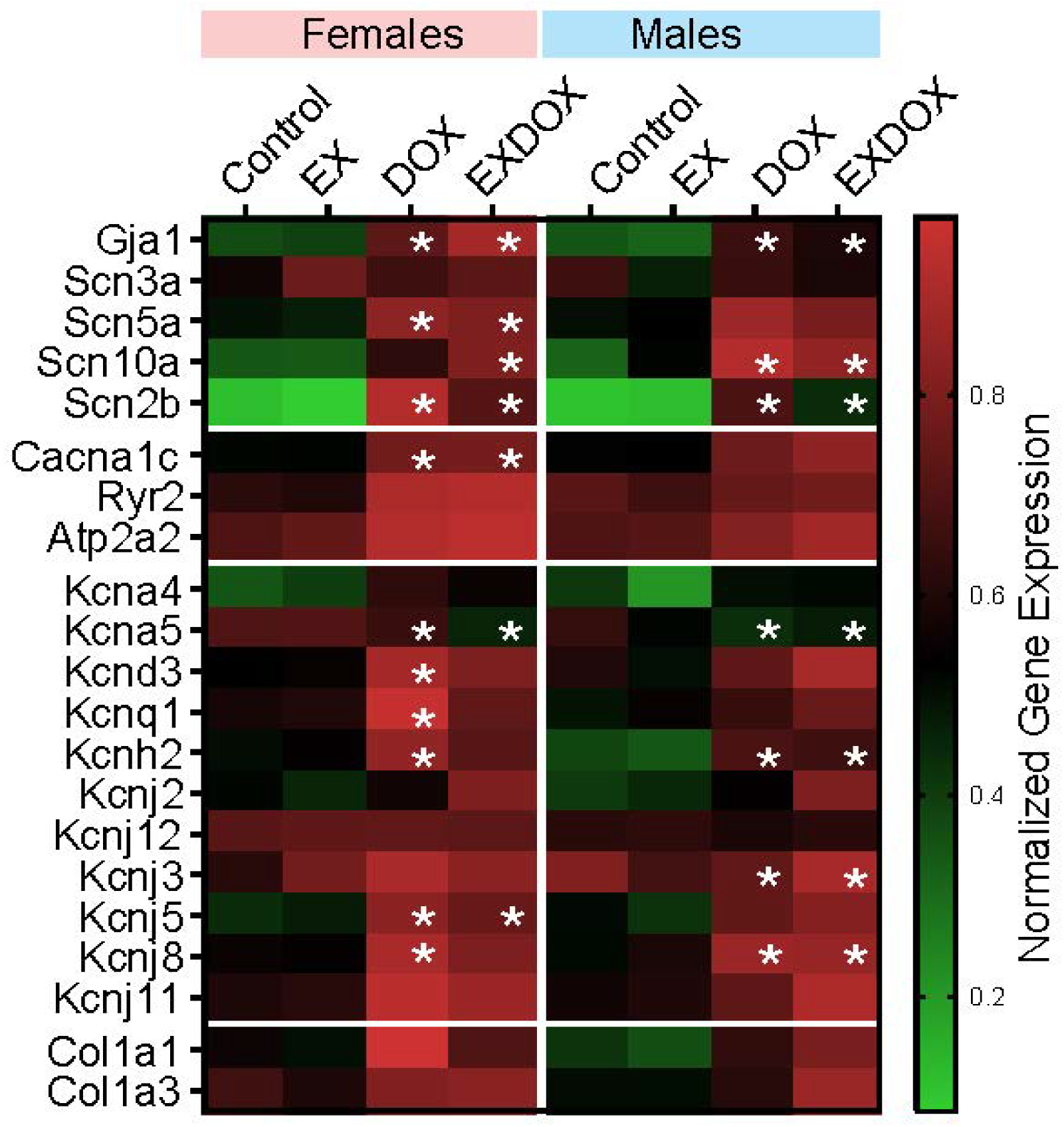
Modulation of electrophysiology gene expression by DOX and exercise. Heat map illustrating modulation of ion channels, gap junctional, calcium handling and fibrosis related genes in male and female hearts from the four experimental groups, Control, EX, DOX and EXDOX. Sample sizes: In each experimental group, 3 biological replicates were used. DEGs were determined using the DESeq2 package in R which uses the Wald test to determine statistical significance and Benjamini-Hochberg method for multiple comparison correction.

## RESULTS

*Exercise delays onset of DOX-induced morbidity:* Total body weight in the EX group did not increase over time as it did in the Control group in both sexes (Figure 1B, Upper Panels). DOX treatment decreased total body weight over the six-week study period, regardless of exercise, in both sexes. Furthermore, DOX reduced health score, indicating increased morbidity in both males and females (Figure 1B, Lower Panels). Importantly, even though morbidity was increased in the EXDOX group compared to Control, it was significantly attenuated compared to DOX group (indicated by # in Figure 1B). Specifically, in EXDOX mice of both sexes, decline in health score was delayed and occurred closer to 5 weeks compared to 2-3 weeks in DOX mice, which could indicate that exercise is able to attenuate DOX induced morbidity until a higher cumulative dose of DOX is achieved (cumulative DOX dose = 25 mg/kg at 5 weeks).

*Exercise fails to prevent DOX-induced mechanical dysfunction in males:* Echocardiographic analysis was performed at the start and end of the study protocol and. Representative echocardiograms from males and females in each experimental group is presented in Figure 2A and normalized summary data is presented in Figure 2B and 2C. Data presented in Figure 2B and 2C are normalized for each mouse at the end of the study protocol compared to its own baseline.

SV was significantly reduced in male DOX and EXDOX groups indicating decline in mechanical function, along with reduced LVEDD. Interestingly, even though LVEDD was reduced in female EXDOX mice, no other echocardiographic parameters were altered in this experimental group. CO was not different in either DOX or EXDOX groups due to a non-significant increase in heart rate (CO = SV x heart rate). Lastly, EF was not altered by DOX or EXDOX in either sex. This finding is in line with our previous studies where we reported both systolic and diastolic dysfunction in our mouse DOX model which led to no change in the EF ratio. A decreased in EF was observed in the female EX group, due to LV dilation (increased LVEDD and LVESD). FS followed the same trend in all groups.

Together, this data suggests 1) DOX altered cardiac mechanical function in males but not in females, and 2) EXDOX failed to prevent these DOX induced effects. *Exercise prevented arrhythmogenic electrical remodeling in female hearts:* Representative optical action potentials and calcium transients from female hearts along with voltage activation maps illustrating the spread of electrical excitation is presented in Figure 3A. Ten parameters of cardiac metabolism-excitation-contraction coupling (MECC) were calculated at varying BCLs and are presented in Figure 3B (females) and 3C (males).

In female DOX mice, APD_80_ prolongation, CV_T_ and CV_L_ slowing and increased Ca τ were observed. Each of these DOX-induced effects could increase risk of arrhythmias. Importantly, in female EXDOX hearts, DOX induced CV_T_ and CV_L_ slowing as well as increase in Ca τ were prevented by exercise, although APD_80_ was still prolonged, particularly at faster pacing rates. Additionally, V_m_RT was significantly reduced, and CaRT was prolonged in female EXDOX hearts indicating shorter depolarization time and longer calcium release times. No significant changes in cardiac MECC parameters were observed in any of the male experimental groups or female EX group.

Taken together with the echocardiographic data above, this indicates a different manifestation of DOX cardiotoxicity in females compared to males. Importantly, while exercise did not prevent DOX induced mechanical dysfunction in males, it attenuated DOX induced electrical remodeling in female hearts.

*Exercise prevented DOX induced metabolic impairment:* OCR which is an index for aerobic metabolism and ECAR which is an index for anaerobic metabolism were calculated by performing seahorse analysis on LV slices from hearts in each group and are presented in Figure 4A. Energy profile plots for male and female groups are illustrated in Figure 4B.

In EX mice, ECAR was increased in both males and females while OCR had an increasing trend (p=0.07) in female hearts alone. On the other hand, DOX treatment reduced OCR and demonstrated a reducing trend (p=0.07) in ECAR in female mice with no significant changes in males. Lastly, OCR and ECAR were preserved in both male and female EXDOX groups and were similar to Controls. This indicates that in female hearts DOX impairs aerobic metabolism, which is the primary source of ATP production, while it is preserved in EXDOX mice. Preserved aerobic metabolism could be a mechanism of cardioprotection in EXDOX females.

*DOX alters cardiac transcriptome and activates inflammatory pathways while exercise is anti-inflammatory:* The PCA plot in Figure 5A illustrates a clear distinction of experimental groups based on DOX administration. Control and EX groups clustered together while DOX and EXDOX groups clustered together indicating that the majority of the changes in the cardiac transcriptome were induced by DOX compared to exercise. Male and female samples did not cluster separately. This is further illustrated in the heatmap generated by unsupervised hierarchical clustering in Figure 5B.

Volcano plots in Figure 5C illustrate that while very few DEGs were observed in the EX group (Females = 26 upregulated DEGs, Males = 2 downregulated DEGs) compared to Controls, DOX and EXDOX groups had a large number of DEGs versus Controls. Specifically, in the female DOX hearts 276 DEGs were upregulated and 768 DEGs were downregulation compared to Control, while in males 154 and 909 DEGs were up- and downregulated, respectively. In the EXDOX group, 374 and 791 DEGs were up- and downregulated in females while 234 and 928 DEGs were up- and downregulated in males, respectively.

Lastly, to determine the specific pathways regulated by the significant DEGs, gene set enrichment analysis (GSEA) was performed. Top ten up- and downregulated pathways whose gene sets were significantly altered by DOX and EXDOX treatments versus Control are illustrated in Figure 5D. In female mice, DOX activated several inflammation-related pathways including TNF signaling pathway and NF-κB signaling pathway. Activation of these pro-inflammatory pathways was prevented in the female EXDOX group. Additionally, in the female DOX group, lipid and atherosclerosis related genes were also enriched which could indicate aberrant lipid metabolism and lipid accumulation. On the other hand, in female EXDOX group, oxidative phosphorylation genes and cardiac muscle contraction genes were enriched. In male mice, calcium signaling genes were downregulated by DOX, but this was prevented in the EXDOX group.

To summarize, while DOX treatment induced a much larger change in the cardiac transcriptome compared to exercise alone, some key differences in enriched gene sets were identified between DOX and EXDOX groups. Of particular note is the enrichment of inflammatory gene sets in the female DOX group, which was prevented in the EXDOX group, indicating that inflammation plays a key role in female DOX cardiotoxicity and exercise induced cardioprotection in this group.

*DOX is pro-inflammatory in females and exercise prevents it:* Next, the protein level expression of 40 cytokines and chemokines were determined in female hearts from the DOX and EXDOX groups (Figure 6A and 6B).

DOX increased the expression of pro-inflammatory molecules such as Eotaxin, interleukin-2 (IL-2), interleukin-17 (IL-17), chemokine (C-X-C motif) ligand 1 (CXCL1), chemokine (C-C motif) ligand 3 (CCL3) and tissue inhibitor of metalloproteinases 1 (TIMP1). EXDOX treatment prevented the DOX-induced increase in IL-17, CCL3 and TIMP-1 but not eotaxin. Futhermore, several other pro-inflammatory cytokines such as chemokine ligand 13 (CXCL13), fractalkaline, interleukin-12-p70 and soluble tumor necrosis factor receptor type II (sTNF RII) were significantly reduced in the EXDOX hearts compared to Control.

DOX also increased the expression of several molecules known to have anti-inflammatory properties including granulocyte colony stimulating factor (G-CSF), interleukin-10 (IL-10), interleukin-12-p40/p70 (IL-12-p40/p70), interleukin-13 (IL-13) and tissue inhibitor of metalloproteinases 2 (TIMP2). EXDOX treatment also increased IL-12-p40/p70, IL-13 and TIMP2 and additionally increased interleukin-4 (IL-4) expression as well.

Considering this protein level data along with the enriched gene set analysis above, it can be concluded that DOX treatment sets up a pro-inflammatory condition in female hearts while EXDOX prevents it.

*DOX and exercise modulate cardiac ion channel gene expression:* Inflammation is known to alter several factors that can affect cardiac electrophysiology and conduction including ion channel and gap junctional protein expression and function, fibrosis, and calcium handling proteins. The modulation of genes associated with these variables by DOX and exercise were next assessed and illustrated in Figure 7. All gene expression values were normalized to the maximum numerical value for that given gene among all experimental groups.

EX did not alter any of the analyzed genes compared to Control. DOX and EXDOX, on the other hand, altered several genes expression. Scn5a, Cacna1c and Kcnj5 were increased in both female DOX and female EXDOX hearts compared to Control, but not in males. On the other hand, Kcnj3 was increased in DOX and EXDOX males versus Control but not females. Additionally, Scn2b, Kcna5 and Gja1 were increased in DOX and EXDOX mice of both sexes versus Control.

Notably, in males, any genes that were modulated by DOX was also similarly altered in the EXDOX group. Contrarily, there were several differences in female DOX (arrhythmogenic electrical remodeling) versus female EXDOX (preserved electrophysiology) groups. Specifically, potassium channel genes Kcnd3, Kcnq1, Kcnh2 and Kcnj8 were increased in female DOX hearts but not in EXDOX while Scn10a was increased in female EXDOX hearts but not in DOX.

Thus, DOX modulates several ion channel gene expressions which could underlie the aberrant electrical function in female DOX treated hearts. Specifically, increase in Cacna1c correlates with APD_80_ prolongation in DOX and EXDOX females while modulation of potassium ion channel genes that set the resting membrane potential in female DOX mice could explain CV slowing in this group.

## DISCUSSION

In this study, we demonstrate that not only is DOX cardiotoxicity sex-specific but that exercise therapy to prevent DOX cardiotoxicity also has sex-specific effects. Specifically, while exercise prevented DOX-induced arrhythmogenic electrical remodeling in female hearts, it did not alter cardiac electrical or mechanical function in DOX treated male hearts. DOX induced aberrations in female cardiac electrophysiology was associated with impaired aerobic metabolism, activation of inflammatory pathways and modulation of ion channel gene expression in the heart. Importantly, exercise therapy restored cardiac metabolism, was anti-inflammatory and prevented several ion channel remodeling during DOX treatment. Therefore, exercise therapy could have significant cardioprotective effects in female cancer patients undergoing DOX therapy but not in males. These findings highlight the critical need for sex-informed therapies in cardio-oncology and cardiovascular diseases.

*Sex-specific DOX cardiotoxicity:* DOX cardiotoxicity is sex-specific.^4^ Female patients are associated with greater risk of adverse cardiovascular effects among pediatric patients, lower risk among premenopausal adults and similar risk among postmenopausal adults compared to age-matched men. DOX cardiotoxicity is not as prevalent among premenopausal adult women compared to similarly aged men and the protective effects of estrogen have been hypothesized to be responsible for cardioprotection among these patients. However, we demonstrate here that while DOX-induced mechanical dysfunction was not observed in female mice, DOX caused significant arrhythmogenic electrical remodeling in these female hearts. On the other hand, DOX treated male mice had impaired cardiac mechanical function but preserved electrophysiology. This finding is in line with our previous reports in acute DOX treated mouse and human heart models where DOX detrimentally modulated electrophysiology in female hearts but not in males.^23,24^ Specifically, in mice, 5 days after a single intraperitoneal injection of DOX, P and QRS durations were reduced and QTc and RR intervals were increased in females but not in males.^23^ Similarly, 24-hour incubation of human cardiac organotypic slices with DOX induced APD prolongation in young (<50 years of age) female hearts but not age-matched males.^24^

Clinically, cardiotoxicity is primarily determined by deterioration in mechanical function and increase in global longitudinal strain. Reduction in EF to less than 50-55% or a >10% decline from a patient’s pre-treatment EF is typically used to define DOX cardiotoxicity.^5,6^ While EKGs are recorded in cancer patients undergoing chemotherapy, due to prevalence of arrhythmias during and immediately after DOX chemotherapy, they are typically not used to define cardiotoxicity. Together with our previous publications, this study highlights the critical need to redefine cardiotoxicity by assessing multiple aspects of cardiac physiology, looking beyond mechanical dysfunction. Using a sex-informed diagnostic approach for cardiotoxicity will prevent female cancer patients from becoming an underdiagnosed population with a higher risk of arrhythmias.

*Exercise therapy modulates cardiac metabolism:* Fatty acids and glucose are the two primary metabolic substrates used to produce ATP that powers several aspects of cardiac physiology including contraction and ion channel function. Aerobic exercise such as running, swimming, and biking improves oxidation of these metabolic substrates and increases ATP production.^25^ Moderate intensity exercise is prescribed for healthy individuals and those with cardiovascular diseases.^26,27^ Exercise improves cardiac function and reduces the severity of several cardiovascular diseases such as myocardial infarction, hypertension, and heart failure.

Modulation of cardiac metabolism and substrate preference vary depending on exercise modality, duration and if the measurement was made during exercise or at rest.^28–31^ In this study, we measured resting metabolism in mice treated with DOX and/or exercise and we demonstrate that exercise modulated aerobic metabolism in female hearts. Oxygen consumption by cardiac tissue, a metric of aerobic metabolism, demonstrated an increasing trend (p = 0.07) in female exercised hearts while DOX treatment significantly reduced it. More importantly, exercise during DOX treatment prevented DOX-induced impairment of aerobic metabolism in these female hearts. Contrarily, in male hearts, exercise improved extracellular acidification, a metric of anaerobic metabolism both with (p=0.07) and without (p<0.05) DOX treatment. The majority of the ATP produced in the heart comes from aerobic oxidation of fatty acids and glucose. The ATP thus produced is used in contraction and in proper functioning of ion channels and pumps, among other physiological processes. By reducing aerobic metabolism in female hearts, DOX could alter ion currents and action potential generation and propagation and increase risk of arrhythmias. On the other hand, only a small percentage of total ATP production is dependent on anaerobic metabolism. Thus, the modulation of anerobic metabolism in male hearts may not be sufficient to affect male cardiac physiology.

*Exercise therapy is anti-inflammatory in DOX treated female hearts:* DOX activates inflammatory pathways and increases expression of inflammatory cytokines.^32–34^ DOX increases TNFα, IL-8, and NF-κB expression which have been implicated in the progression of cardiotoxicity and congestive heart failure.^35^ In this study, we demonstrate in a six-week DOX treated mouse model that DOX increased the mRNA and protein level expression of several inflammatory markers in female hearts, but not in males. This is the first study to report a sex-specific inflammatory response to DOX treatment. Specifically, genes involved in the pro-inflammatory TNFα and NF-κB signaling were increased in DOX treated female hearts and protein expression of pro-inflammatory cytokines - eotaxin, IL-2, IL-17, Cxcl1, Ccl3 and TIMP-1 were also increased. Interestingly, TNFα and its receptor (TNFR1) were not increased in expression by DOX, either at the mRNA or protein level, even though the TNFR1-associated death domain (TRADD) gene and its downstream targets were increased. TRADD binds to TNFR1 and transduces the downstream signaling pathways involved in apoptosis, proliferation and NF-κB activation.^35^ Alternative ligands that can activate TRADD and the TNFα signaling pathway are currently unknown.

Moderate levels of regular exercise has known anti-inflammatory effects via the reduction in visceral fat mass,^36,37^ decrease in adipokines and pro-inflammatory cytokines,^38,39^ and reduced expression of toll-like receptors on immune cells.^40,41^ Increased expression of anti-inflammatory cytokines such as IL-6 and IL-10 expression increase have also been associated with the protective effects of exercise in chronic inflammatory conditions.^42,43^ In this study, DOX induced a chronic pro-inflammatory condition in female mice and consistent with previous reports,^36^ exercise attenuated inflammation in the female EXDOX mice. Exercise prevented the DOX-induced increase in pro-inflammatory cytokines and chemokines such as IL-17, Ccl3 and TIMP-1, but failed to modulate others such as IL-2, IL-13 and TIMP-2. Additionally, gene set enrichment analysis revealed that exercise prevented the DOX-induced activation of the TNFα and NF-κB signaling pathways. Together, these data indicate that in this chronic DOX treatment model, DOX induced a pro-inflammatory condition in a sex-specific manner while exercise therapy attenuated it.

*Ionic basis of exercise induced cardioprotection in female hearts:* Metabolic impairment and inflammation can both alter ion channel expression and function to promote an arrhythmogenic substrate.^44–47^ Here, we report that DOX reduced aerobic metabolism and activated inflammatory cytokines in female hearts but not males. These changes correlated with modulation of ion channel gene expression including increase in Cacna1c, Scn5a and several genes that encode for potassium channels in female hearts but not males. The functional consequence of DOX induced modulation of these ion channels was APD prolongation and CV slowing in DOX treated female hearts.

Consistent with the findings of this study, we previously reported that acute DOX treatment induced APD prolongation in young female human donor heart slices, but not males, and this correlated with a similar increase in Cacna1c which encodes for Ca_v_1.2 responsible for the L-type calcium current (I_CaL_).^24^ Other studies have also reported an increase in I_CaL_ in response to DOX treatment.^48–50^ Early repolarization is a fine balance between opposing calcium and potassium currents and an increase in one can alter the morphology and duration of the action potential. Thus, under any given condition, the relative contribution of each of these ion currents controls APD. A previous study reported that the increase in I_CaL_ was a principle determinant of the effects of acute and chronic DOX treatment on APD.^51^ Modulation of I_Kr_ (increased by acute DOX and decreased by chronic DOX) also affected DOX-induced APD modulation.^51^ Thus, upregulated Cacna1c and therefore increased I_CaL_ could underlie the female sex-specific APD prolongation by DOX. Lastly, exercise did not prevent the DOX-induced increase in Cacna1c and APD prolongation.

DOX also slowed conduction velocity in female hearts. Conduction velocity is modulated by tissue excitability, intercellular coupling and tissue architecture which includes cellular geometry and extracellular spaces. The increase in Kcnj8 which encodes for K_ir_6.1 responsible for the ATP-sensitive inward rectifier potassium current (I_KATP_) could affect tissue excitability. I_KATP_ is activated in response to decrease in ATP and increase in ADP induced by metabolic impairment and hyperpolarizes the cell which in turn reduces cellular excitability. Thus, DOX-induced CV slowing in female hearts could be an effect of increased Kcnj8 expression and metabolic impairment. Importantly, we also show here that exercise, by preventing DOX-induced aerobic metabolic impairment and Kcnj8 upregulation, preserves cardiac conduction.

DOX also modulates other factors that can affect conduction velocity. DOX increased Gja1 and Scn5a which encode for the gap junctional protein, Cx43, and the voltage gated sodium channel responsible for the sodium current, I_Na_. Typically, decrease in these proteins which reduces intercellular coupling, is associated with conduction slowing.^52–54^ However, the intercellular coupling – conduction velocity relationship is further modulated by extracellular volumes in nanodomains along the intercalated discs.^54^ In male hearts, DOX increased the width and area of the perinexus, a site critical for ephaptic intercellular coupling, but not in females (data not shown). Slight increases of this extracellular nanodomain, in the setting of disease conditions such as metabolic ischemia and hyponatremia preserves conduction while narrowing of this space causes conduction slowing by self-attenuation of sodium channels.^55,56^ Thus, DOX-induced widening of the perinexal space in males could underlie preserved conduction while the lack of this response in female hearts could also contribute to conduction slowing.

*Limitations:* In this chronic study, while we assessed progression of mechanical function modulation by DOX, we only measured electrophysiology, metabolism and underlying mechanistic pathways at the end of the six-week study period. This limits us from understanding the causative role of each of the tested factors in DOX cardiotoxicity.

## ACKNOWLEDGEMENTS

We gratefully acknowledge Drs Krithika Rao and Sruti Shiva from the Center for Metabolism and Mitochondrial Medicine (C3M) at the University of Pittsburgh for their support with the seahorse metabolic analysis. We also thank Dr Igor Efimov for his guidance and Bhavi Barnwal for her support in this project.

## SOURCES OF FUNDING

This study was supported by an American Heart Association Career Development Award (935807) and Bridge Career Development Award (26CBDA1622701) to SAG as well as startup funds from the University of Pittsburgh to SAG.

## DISCLOSURES

None.

## NON-STANDARD ABBREVIATIONS AND ACRONYMS

EX: exercised mouse group
DOX: doxorubicin treated mouse group
EXDOX: exercise + doxorubicin treated mouse group
LV: left ventricle
EF: ejection fraction
FS: fractional shortening
SV: stroke volume
CO: cardiac output
LVEDD: left ventricular end diastolic diameter
LVESD: left ventricular end systolic diameter
BCL: basic cycle length
NADH: nicotinamide adenine dinucleotide hydride
VmRT: voltage rise time
APD80: action potential duration at 80% repolarization
CVT: transverse conduction velocity
CVL: longitudinal conduction velocity
AR: anisotropic ratio
CaRT: calcium rise time
CaTD80: calcium transient duration at 80% calcium reuptake
Ca τ: calcium decay constant
Vm-Ca: delay voltage to calcium activation delay
OCR: oxygen consumption rate
ECAR: extracellular acidification rate
DEG: differentially expressed genes
PCA: principal component analysis
GSEA: gene set enrichment analysis

## Notes

### Competing Interest Statement

The authors have declared no competing interest.

